# Cerebellar-Prefrontal Connectivity Predicts Negative Symptom Severity Across the Psychosis Spectrum

**DOI:** 10.1101/2024.11.07.622549

**Authors:** Sean A. Yarrell, Sophia H. Blyth, Baxter P. Rogers, Anna Huang, Alexandra B. Moussa-Tooks, Neil D. Woodward, Stephan Heckers, Roscoe O. Brady, Heather Burrell Ward

## Abstract

**Background:** Negative symptom severity predicts functional outcome and quality life in people with psychosis. However, negative symptoms are poorly responsive to antipsychotic medication and existing literature has not converged on their neurobiological basis. Previous work in small schizophrenia samples has observed that lower cerebellar-prefrontal connectivity is associated with higher negative symptom severity and demonstrated in a separate neuromodulation experiment that increasing cerebellar-prefrontal connectivity reduced negative symptom severity. We sought to expand this finding to test associations between cerebellar-prefrontal connectivity with negative symptom severity and cognitive performance in a large, transdiagnostic sample of individuals with psychotic disorders.

**Methods:** In this study, 260 individuals with psychotic disorders underwent resting-state MRI and clinical characterization. Negative symptom severity was measured using the Positive and Negative Symptoms Scale, and cognitive performance was assessed with the Screen for Cognitive Impairment in Psychiatry. Using a previously identified cerebellar region as a seed, we performed seed to whole brain analyses and regressed connectivity against negative symptom severity, using age and sex as covariates.

**Results:** Consistent with prior work, we identified relationships between higher cerebellar-prefrontal connectivity and lower negative symptom severity (r=-0.17, p=.007). Higher cerebellar-prefrontal connectivity was also associated with better delayed verbal learning (r=.13, p=.034).

**Conclusions:** Our results provide further evidence supporting the relationship between cerebellar-prefrontal connectivity and negative symptom severity and cognitive performance. Larger, randomized, sham-controlled neuromodulation studies should test if increasing cerebellar-prefrontal connectivity leads to reductions in negative symptoms in psychosis.

## Introduction

Schizophrenia affects approximately 24 million people worldwide and is characterized by positive and negative symptoms and cognitive impairment (1). Negative symptoms are defined by emotional, social, and psychomotor impairment (2). This includes blunted affect, alogia, anhedonia, asociality, and avolition (3). Negative symptom severity strongly predicts functional outcomes and overall quality of life (4-6). Critically, negative symptoms are often minimally responsive to antipsychotic medication (7, 8).

To develop more effective treatments for negative symptoms, previous studies have used neuroimaging to investigate the neurobiological basis of negative symptoms. However, existing correlational neuroimaging studies have not yielded consistent findings (9, 10). Neuromodulation interventions, such as repetitive transcranial magnetic stimulation (rTMS) offer the opportunity to test causal associations between brain circuitry and negative symptom severity.

Brady et al. used this approach in 2019, where they combined a novel, data-driven connectome wide multivariate analysis to identify neuroimaging correlates of negative symptom severity in psychotic disorders with an rTMS intervention to test this relationship. In the first part of their analysis, they observed that lower cerebellar-prefrontal connectivity was associated with higher negative symptom severity in 44 individuals with schizophrenia. Then, they applied intermittent theta burst stimulation (iTBS), a form of rTMS associated with excitatory effects, to the cerebellum in 11 individuals with schizophrenia. Compared to sham, in the active stimulation group, increased cerebellar-prefrontal connectivity was associated with greater reductions in negative symptom severity.

While this approach has several strengths, including the use of a neuromodulation intervention to probe brain-behavior relationships in an independent sample, the study relied on small samples (n=44 and n=11) of individuals with schizophrenia or schizoaffective disorder from a single site and did not assess cognitive performance.

We therefore sought to test if cerebellar-prefrontal connectivity was associated with negative symptom severity across the psychosis spectrum using a large (n=260), transdiagnostic sample of individuals with psychotic disorders collected at an independent site. We also extended previous findings by identifying novel relationships between cerebellar-prefrontal connectivity, negative symptom severity, and cognitive performance.

## Methods

### Participants

Data came from a repository of 361 individuals with psychotic disorders with complete neuroimaging and behavioral data who participated in one of three neuroimaging projects conducted in the Department of Psychiatry and Behavioral Sciences at Vanderbilt University Medical Center (VUMC, CT00762866; 1R01MH070560; 1R01MH102266). Participants were recruited from the Psychotic Disorders Program at VUMC. All studies were approved by the Vanderbilt Institutional Review Board (IRB) and all individuals provided written informed consent prior to participating in the studies. The Structured Clinical Interview for DSM-IV or DSM-5 (SCID-IV, SCID-5) was administered to all study participants to confirm diagnoses (11, 12). Exclusion criteria were similar across the three studies, including age under 16/18 years or over 65 years (55 years in 1R01MH102266); estimated premorbid IQ less than 70 based on the Wechsler Test of Adult Reading; history of significant head trauma, medical illness, or central nervous system disorder; pregnancy or lactation; substance abuse within the last 1 month for patients (3 months in 1R01MH102266); and MRI contraindicators. After quality control (see *MRI Data Processing*), the final sample consisted of 260 individuals with a psychotic disorder (schizophrenia n=81; schizophreniform n=73; bipolar disorder with psychotic features n=71; schizoaffective disorder n=28; brief psychotic disorder n=3, major depressive disorder with psychotic features n=3; other psychotic disorder n=1, Table 1).

**Table 1.** Demographics.

| **Demographic** | **Psychosis (n=260)** |
| --- | --- |
| Female, n (%) | 87 (33.5) |
| Age, Mean (SD) | 28.2 (11.0) |
| Race, n (%) |  |
| White | 180 (69.2) |
| Black | 66 (25.4) |
| Asian | 2 (0.77) |
| Native American | 3 (1.6) |
| Other | 9 (3.5) |
| PANSS, Mean (SD) |  |
| Positive | 16.1 (8.1) |
| Negative | 13.9 (6.5) |
| General | 29.7 (8.6) |
| Total | 59.7 (18.4) |
| MADRS, Mean (SD) | 9.0 (8.4) |
| SCIP Total Z-Score, Mean (SD) | -0.89 (0.98) |
| Chlorpromazine Equivalents, Mean (SD) | 378.8 (576.6) |
| Diagnosis, n, (%) |  |
| Schizophrenia | 81 (31.2) |
| Schizophreniform | 73 (28.1) |
| Bipolar Disorder with Psychotic Features | 71 (27.3) |
| Schizoaffective Disorder | 28 (10.8) |
| Major Depressive Disorder with Psychotic Features | 3 (1.2) |
| Brief Psychotic Disorder | 3 (1.2) |
| Other Psychotic Disorder | 1 (0.38) |

### Assessments

Psychosis symptoms were measured using the Positive and Negative Syndrome Scale (PANSS) (13), which is a clinician-rated assessment of positive, negative, and general psychopathology symptoms over the past two weeks. The Marder factor analysis was used to calculate positive, negative, and general psychopathology subscores (14). The Marder factor of negative symptoms, comprises 7 items of the PANSS: blunted affect, emotional withdrawal, poor rapport, passive/apathetic withdrawal, motor retardation, active social avoidance, and lack of spontaneity in conversation. Depressive symptom severity was assessed using the Montgomery–Åsberg Depression Rating Scale (15).

Cognitive performance was assessed using the Wechsler Test for Adult Reading (WTAR)(16) and the Screen for Cognitive Impairment in Psychiatry (SCIP) (17). The SCIP includes measures of verbal memory (immediate and delayed), working memory, verbal fluency, and processing speed and is a reliable and validated measure of cognitive impairment in psychotic disorders (17, 18). SCIP subtest raw scores were converted to z-scores using normative data and averaged to create a composite z-score.

### MRI Acquisition

Neuroimaging data were collected on one of two identical 3.0-T Philips Intera Achieva MRI scanners (Philips Healthcare, Andover, MA), located at the Vanderbilt University Institute of Imaging Science. Briefly, a seven or ten-minute echo-planar imaging (EPI) resting-state fMRI scan and T1-weighted anatomical (1 mm isotropic resolution) were collected for each participant. Preprocessing of fMRI data, performed in SPM12, included correction for head motion using rigid body motion correction, spatial co-registration to T1-weighted anatomical images, and spatial normalization to MNI-space using the parameters obtained from the grey matter segmentation normalization.

### MRI Data Processing

Anatomical images were segmented into gray matter, white matter and cerebrospinal fluid (CSF) with the Computational Anatomy Toolbox 12 (CAT12, version 12.5; http://www.neuro.uni-jena.de/cat/). Resting-state scans were preprocessed in SPM12 and were (1) realigned to a mean scan, (2) coregistered with the native space structural scan, then (3) underwent resting-state denoising procedures: bandpass filter (0.01–0.1 Hz), regression of CSF and white matter signal, regression of 12 motion parameters (6 translation and rotation parameters and their first derivative). All resting state scans went through a quality assurance procedure that included calculating framewise displacement (FD) and temporal signal to noise ratio (tSNR). Scans with a mean FD > 0.5 (n=36) or a tSNR lower than the 5th percentile of the distribution of the entire sample (n=32) were excluded from further analysis. To ensure adequate coverage of the cerebellum, we excluded scans with <50% coverage of our cerebellar cluster (n=33). After quality control analyses, there were a total of 260 scans for analysis.

### MRI Data Analysis

We calculated cerebellar-prefrontal connectivity by extracting the time course of the blood-oxygen-level-dependent (BOLD) signal from the cerebellar region identified in Brady et al. (19). Using SPM12 (SPM – Statistical Parametric Mapping, http://www.fil.ion.ucl.ac.uk/spm) we regressed the z-transformed Pearson’s correlation coefficient connectivity maps against the PANSS negative symptom subscore, using age and sex as covariates, to generate spatial maps of how whole functional brain connectivity to the cerebellar region varied with negative symptom severity at a threshold of p<.05. To quantify the relationship between cerebellar-prefrontal connectivity with behavioral variables, we then measured region to seed connectivity at this step by measuring BOLD correlation between the Brady et al. cerebellar region and a 6mm sphere (seed) placed at the location of maximal connectivity-negative symptom association that we observed in the left dorsolateral prefrontal cortex (L DLPFC, MNI x = -26, y = +32, z = +38). We then correlated connectivity between the cerebellar region and the seed placed at maximal connectivity-negative symptom association in the L DLPFC with PANSS negative symptom subscore.

### Statistical Analysis

Pearson’s correlation coefficients were used to determine the relationships between functional connectivity and behavioral measures. T-tests were used to compare continuous outcomes based on dichotomous variables. ANOVAs were used to compare continuous outcomes based on three or more groups. All analyses were conducted in RStudio (Version 2023.03.1+446) using alpha<.05.

Using the *mediation* package in R, we conducted a mediation analysis to test whether the relationship between negative symptoms and cerebellar-prefrontal connectivity is mediated by cognitive performance. In the model, the sample consisted of 240 participants. Mediation analysis was conducted using nonparametric bootstrap confidence intervals based on the percentile method and with 1,000 simulations. In the model, the independent variable was negative symptoms, the mediator was delayed verbal learning Z-score, and the dependent variable was cerebellar-prefrontal connectivity.

## Results

### Negative Symptom Severity, but not Cerebellar-Prefrontal Connectivity, is Associated with Psychotic Disorder Diagnosis

In our final sample of 260 individuals with psychosis spectrum disorders, there was a significant association between PANSS negative symptom severity and diagnosis (F=10.902, df=6, p=8.54e-11, Figure 1). Cerebellar-prefrontal connectivity was not associated with diagnosis (F=0.2013, df=6, p=.976, Supplemental Figure 1).

**Figure 1.**
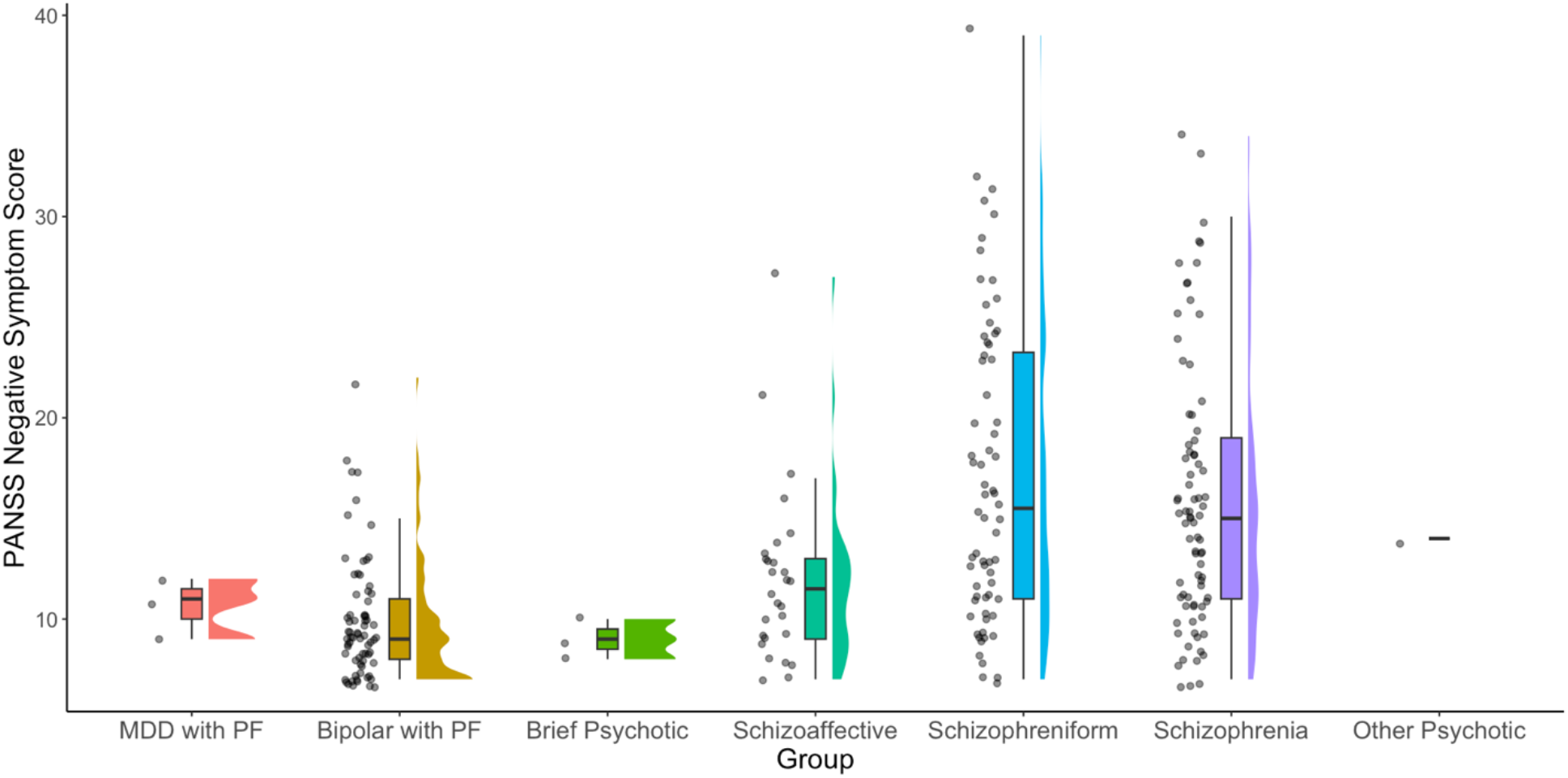
Negative Symptom Severity is Associated with Psychotic Disorder Diagnosis. In a transdiagnostic sample of individuals with psychotic disorders (n=260), PANSS negative symptom severity was associated with diagnosis (F=10.902, df=6, p=8.54e-11).

### Cerebellar-Prefrontal Connectivity is Associated with Negative Symptom Severity Across the Psychosis Spectrum

We observed that higher cerebellar-prefrontal connectivity was associated with lower negative symptom severity (Figure 2A & 2B, r =-0.17, p=.007). This connectivity relationship was specific to negative symptoms, as cerebellar-prefrontal connectivity was not associated with PANSS positive symptoms (r=-0.089, p=.15, Supplemental Figure 3A) or MADRS depressive symptoms (r=-0.040, p=.53, Supplemental Figure 3B).

**Figure 2.**
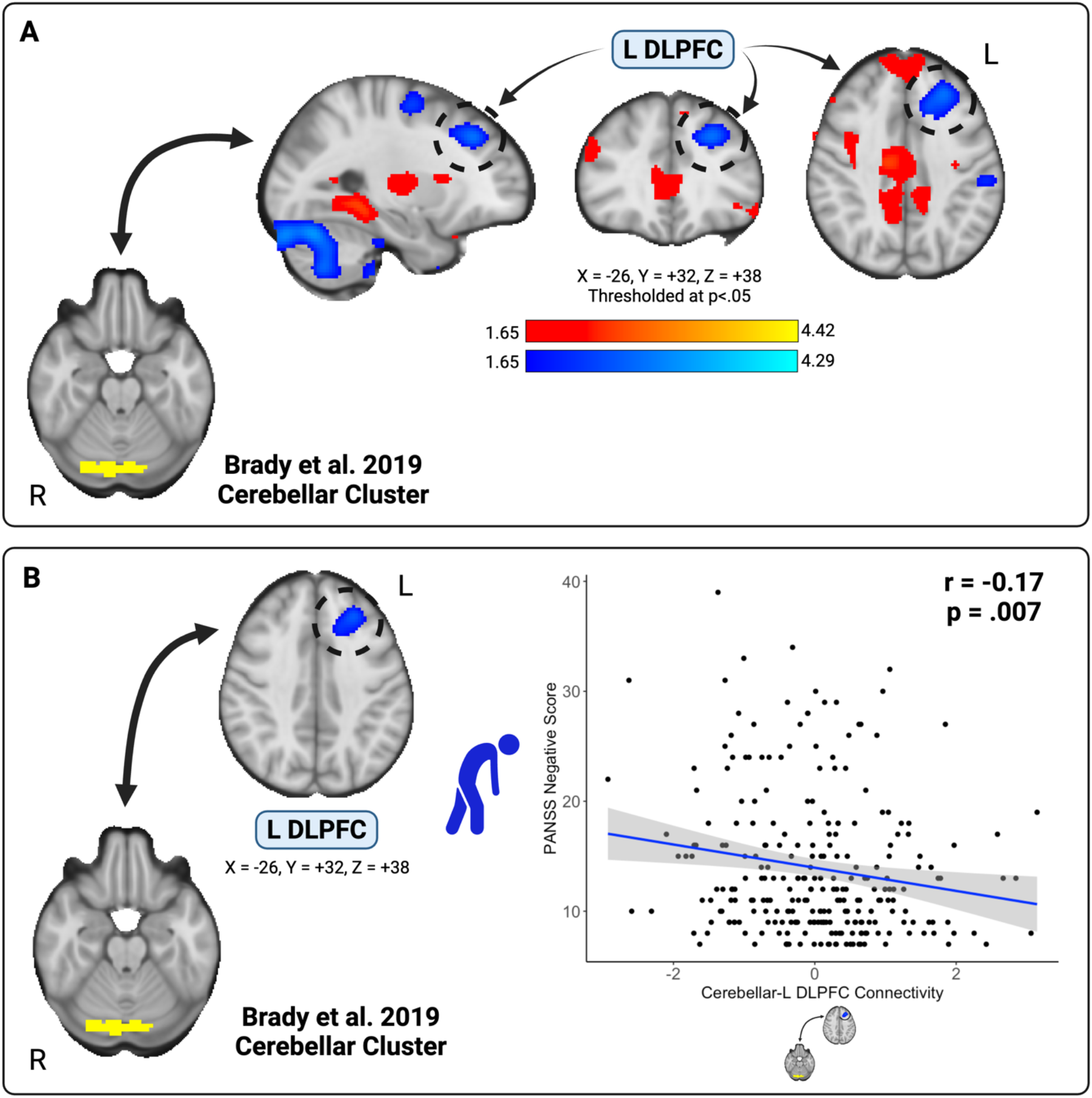
Cerebellar-Prefrontal Connectivity is Associated with Negative Symptom Severity Across the Psychosis Spectrum. We observed that higher cerebellar-prefrontal connectivity was associated with lower negative symptom severity (Figure 2A & 2B, r =-0.17, p=.007). This connectivity relationship was specific to negative symptoms, as cerebellar-prefrontal connectivity was not associated with PANSS positive symptoms (r=-0.089, p=.15) or MADRS depressive symptoms (r=-0.040, p=.53).

To confirm our result was not affected by scanner coverage of our cerebellar region, we repeated our neuroimaging analyses for scans with >70% coverage of the cerebellar region (n=223) and observed similar results such that higher cerebellar-prefrontal connectivity was associated with lower negative symptom severity (Supplemental Figure 2A & 2B, r=-0.18, p=.008). To ensure our result was transdiagnostic and not specific to schizophrenia spectrum disorders, we repeated our analysis in diagnostic subgroups: schizophrenia spectrum disorders (n=182, r=-0.22, p=.006, Supplemental Figure 2C); schizophrenia (n=81, r=-0.11, p=.31, Supplemental Figure 2D); and bipolar disorder with psychotic features (n=71, r=-0.26, p=.027, Supplemental Figure 2E). In all subgroup analyses, we observed similar relationships between cerebellar-prefrontal connectivity and negative symptom severity.

In a general linear model predicting negative symptom severity that included cerebellar-prefrontal connectivity, diagnosis, age, and sex as predictors, cerebellar-prefrontal connectivity and diagnoses of schizophrenia and schizophreniform were significantly related to negative symptom severity (F=8.584, df1=9, df2=250, p=3.373e-11; cerebellar-prefrontal connectivity t=-2.761, p=.0062; schizophrenia t=6.319, p=1.20e-9; schizophreniform t=6.141, p=3.20e-9).

Confirming our results were not affected by antipsychotic medication, chlorpromazine equivalents were not associated with negative symptom severity (r=0.034, p=.63) or cerebellar-prefrontal connectivity (r=-0.062, p=.37).

### Negative Symptoms are Broadly Associated with Cognitive Performance

PANSS negative symptom severity was associated with the WTAR (r=-0.33, p= 7.446e-08) and all cognitive domains on the SCIP: verbal learning – immediate (r=-0.31, p=4.632e-7), working memory (r=-0.34, p=2.063e-8), verbal fluency (r=-0.17, p=.0053), verbal learning-delayed (r=-0.27, p=1.19e-5), processing speed (r=-0.25, p=4.75e-5), and total score (r=-0.38, p=4.58e-10).

### Cerebellar-Prefrontal Connectivity is Associated with Delayed Verbal Learning Across the Psychosis Spectrum

Given existing evidence supporting the role of both the cerebellum and DLPFC in cognitive performance, we investigated the relationships between cerebellar-prefrontal connectivity and cognitive performance. Higher cerebellar-prefrontal connectivity was associated with better delayed verbal learning on the SCIP (r=0.13, p=.034, Figure 3). Connectivity was not related to premorbid IQ or cognitive performance in any other domains of the SCIP.

**Figure 3.**
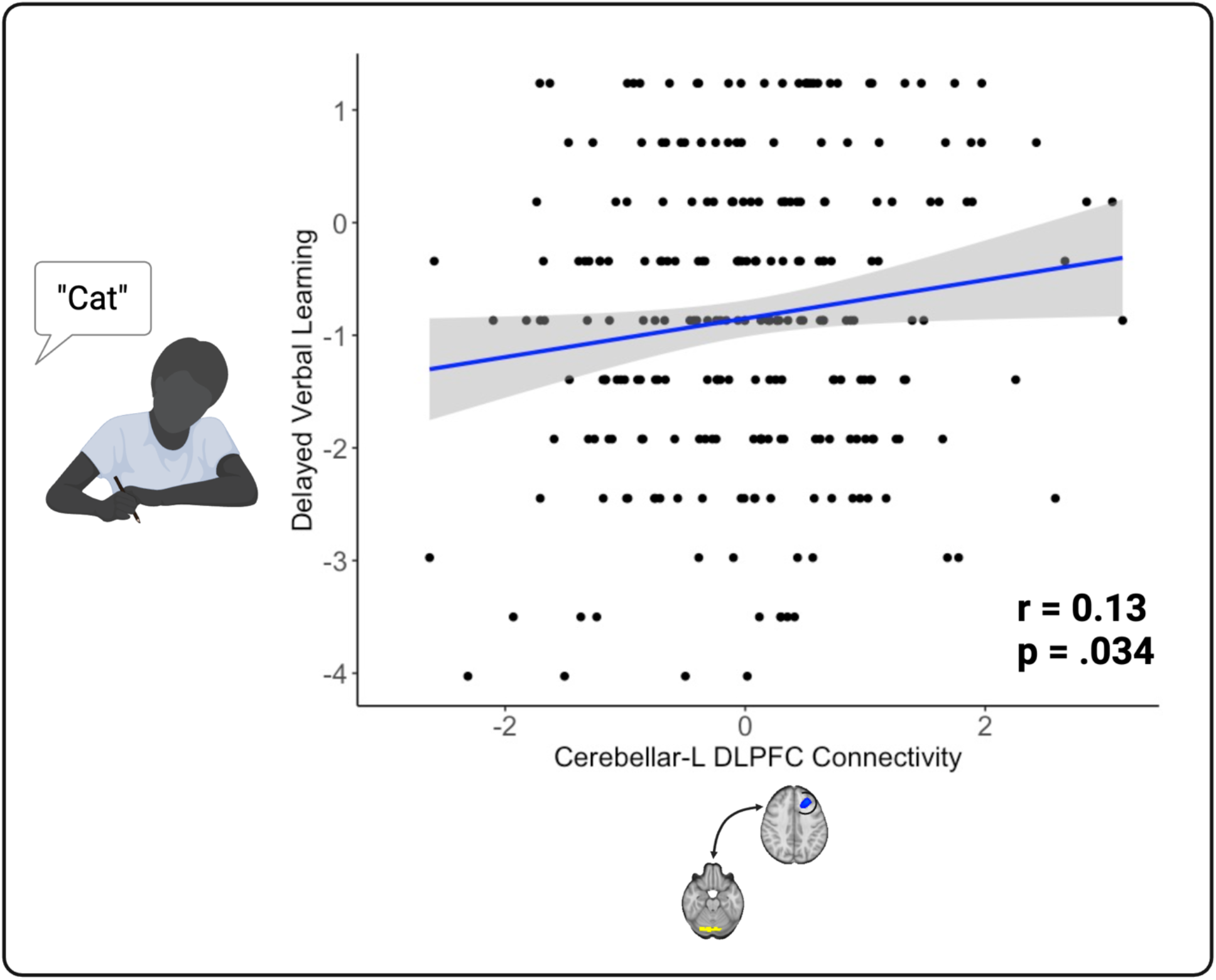
Cerebellar-Prefrontal Connectivity is Associated with Delayed Verbal Learning. Given existing evidence supporting the role of both the cerebellum and DLPFC in cognitive performance, we investigated the relationships between cerebellar-prefrontal connectivity and cognitive performance. Higher cerebellar-prefrontal connectivity was associated with better delayed verbal learning (r=0.13, p=.034). Connectivity was not related to cognitive performance in any other domains.

### Cerebellar-Prefrontal Connectivity Does Not Mediate the Relationship Between Negative Symptom Severity and Delayed Verbal Learning

Given the associations between cerebellar-prefrontal connectivity and both negative symptom severity and delayed verbal learning, we performed a mediation analysis to determine if delayed verbal learning mediated the relationship between cerebellar-prefrontal connectivity and negative symptoms. In this model, the mediation effect of delayed verbal learning Z-score (ACME=-0.2487 [-0.5447, -0.03]) and the proportion of the relationship mediated by delayed verbal learning Z-score (estimate=0.2544 [0.0299-0.78]) were both significant (p=0.030 and p=0.038, respectively). The average direct effect of cerebellar-prefrontal connectivity on negative symptoms (ADE=-0.7289 [-1.4710, -0.06] and the total effect of cerebellar-prefrontal connectivity on negative symptoms (estimate=-0.9776 [-1.7857, -0.26]) were also both significant (p=0.030 and p=0.008, respectively).

These results suggest that there is a significant total and direct effect of cerebellar-prefrontal connectivity on negative symptoms. Additionally, the model suggests that there is a mediating effect of delayed verbal learning Z-score on the relationship between cerebellar prefrontal-connectivity and negative symptoms.

## Discussion

In this study, we sought to extend recent findings linking cerebellar-prefrontal connectivity to negative symptom severity in schizophrenia. Consistent with prior work (19), we observed that higher cerebellar-prefrontal connectivity was associated with lower negative symptom severity.

Our work extends the relationship between cerebellar-prefrontal connectivity and negative symptoms into a 5-fold larger, transdiagnostic sample of psychotic disorders that was collected in an entirely different geographic location. Our findings are also novel because we used the cerebellar region as a seed, whereas Brady et al. used the DLPFC region, and still observed significant relationships to DLPFC regions.

We also observed new relationships between cerebellar-prefrontal connectivity and delayed verbal learning. Cerebellar-prefrontal connectivity was associated with both negative symptoms and delayed verbal learning. Delayed verbal learning mediated the relationship between cerebellar-prefrontal connectivity and negative symptoms. This observation is consistent with previous findings that processing speed mediated the relationship between cerebellar grey matter volume and psychomotor disturbance in psychotic spectrum disorders (20).

Our findings provide additional evidence supporting the role of the cerebellum in social function and cognitive performance. The cerebellum is the most neuron-dense region of the brain; containing roughly 80% of the brain’s neurons (21). While the cerebellum has most widely been associated with balance and coordination, recent research has demonstrated the cerebellum’s role in social function and cognitive performance. The cerebellar region we studied has been linked to social, linguistic, and spatial function in a recent precision functional mapping analysis (22), consistent with our findings relating connectivity from this region to negative symptoms and verbal learning.

The relationships we identified between cerebellar connectivity and negative symptoms provide additional evidence that the cerebellum may be a central driver of multiple domains of psychiatric symptoms and cognitive performance. Another recent analysis by our group linked cerebellar connectivity to the cognitive and motor components of psychomotor performance (23). Negative symptoms are a heterogeneous construct consisting of decreased speech production, amotivation, anhedonia, and asociality. Therefore, it is likely that the cerebellum has a common role in negative symptoms and psychomotor function.

Our results also provide further support for cerebellar neuromodulation interventions. Although small, the cerebellar rTMS study from Brady and colleagues demonstrated that increasing cerebellar-prefrontal connectivity led to reductions in negative symptom severity. Given this network edge has nodes in the cerebellum and DLPFC, it is possible that rTMS interventions could use either region as a target. However, given the neuronal density of the cerebellum and recent work showing dramatic individual differences in network topography of the prefrontal cortex (24), the cerebellum may be a preferable rTMS target. Cerebellar rTMS has been extensively studied in neurologic and motor disorders (25-27) and is now being increasingly studied in schizophrenia (28).

Strengths of our study include the use of a large, transdiagnostic sample of individuals with psychotic disorders across all phases of illness, including both early and chronic illness. Participants were taking a range of antipsychotic medications, which were not associated with negative symptoms or cerebellar-prefrontal connectivity. Finally, our sensitivity analyses ensured that the relationships we observed between connectivity and negative symptoms were truly transdiagnostic.

The primary limitation of our study is that our sample was cross-sectional from one site, albeit an entirely different geographic location than the study from Brady and colleagues.

In conclusion, we have used data from a large, transdiagnostic sample of individuals with psychotic disorders to provide further evidence supporting the association between cerebellar-prefrontal connectivity with negative symptom severity. These results should be tested in a larger, randomized, sham-controlled trial of cerebellar rTMS in people with psychotic disorders.

## Supplemental Material

**Supplemental Figure 1.**
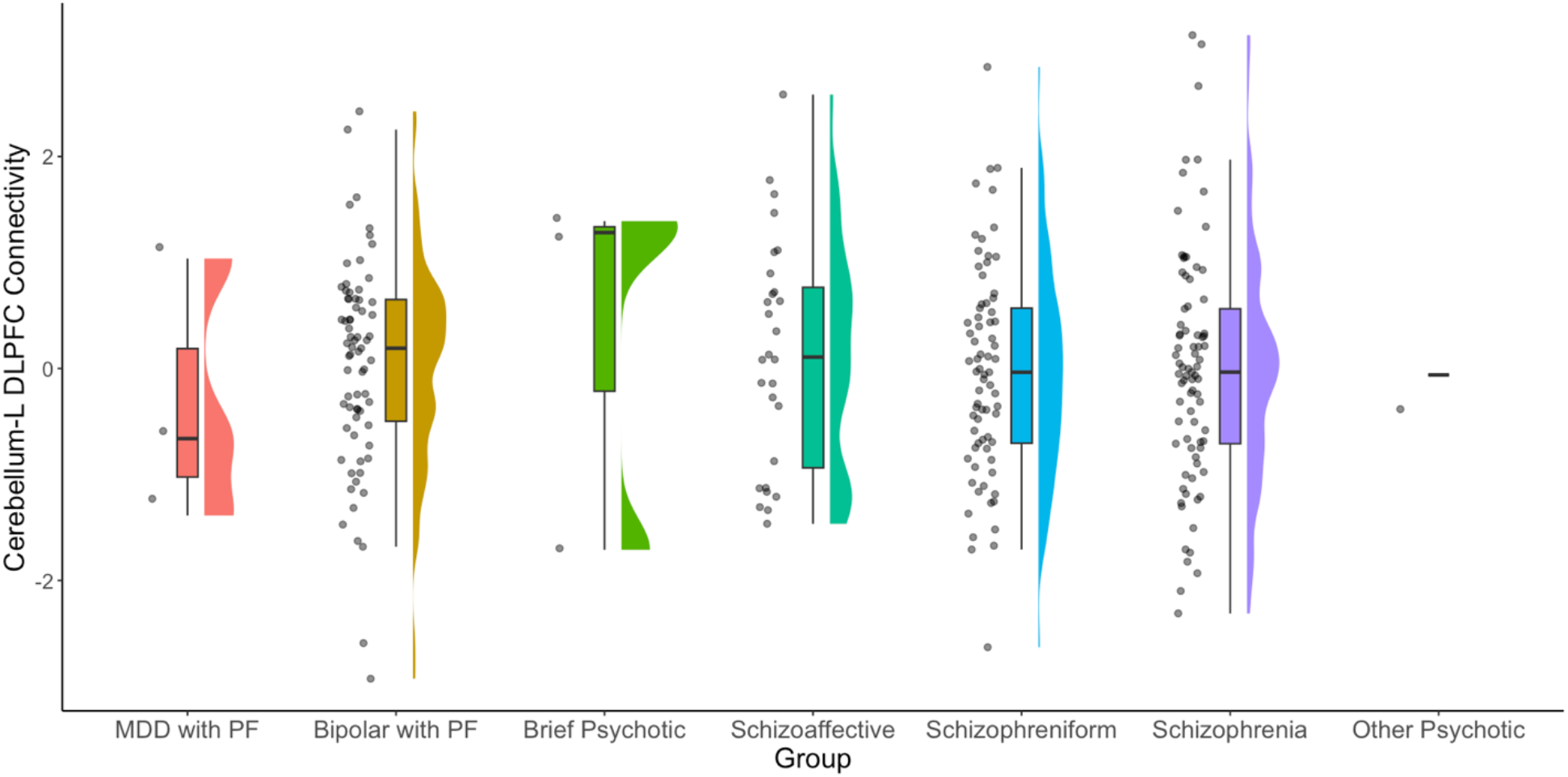
Cerebellar-Prefrontal Connectivity Is Not Associated with Psychotic Disorder Diagnosis. In our sample of 260 individuals with psychotic disorders, we calculated connectivity between the previously identified cerebellar region from Brady and colleagues (19) and the left dorsolateral prefrontal cortex (L DLFPC) region we identified in our analyses. Cerebellar-prefrontal connectivity did not differ by psychotic disorder diagnosis (F=0.2013, df=6, p=.976).

**Supplemental Figure 2.**
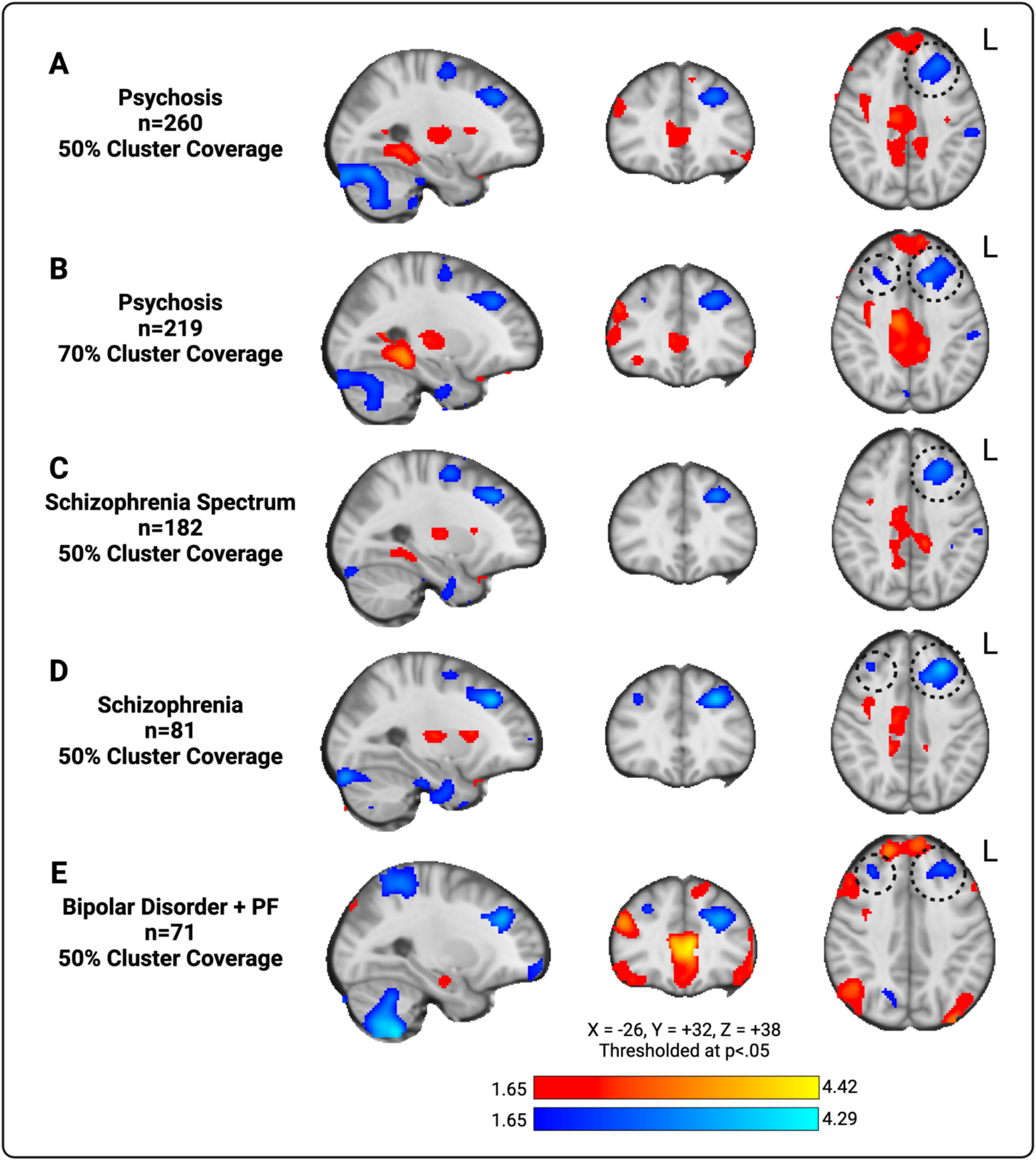
Sensitivity Analyses: Cerebellar-Prefrontal Connectivity is Associated with Negative Symptom Severity Across the Psychosis Spectrum. To confirm our result was not affected by scanner coverage of our cerebellar region, we repeated our neuroimaging analyses for scans with >70% coverage of the cerebellar region (n=223) and observed similar results such that higher cerebellar-prefrontal connectivity was associated with lower negative symptom severity (Supplemental Figure 2A & 2B, r=-0.18, p=.008). To ensure our result was transdiagnostic and not specific to schizophrenia spectrum disorders, we repeated our analysis in diagnostic subgroups: schizophrenia spectrum disorders (n=182, r=-0.18, p=.018, Supplemental Figure 2C); schizophrenia (n=81, r=-0.11, p=.31, Supplemental Figure 2D); and bipolar disorder with psychotic features (n=71, r=-0.26, p=.027, Supplemental Figure 2E). In all subgroup analyses, we observed similar relationships between cerebellar-prefrontal connectivity and negative symptom severity.

**Supplemental Figure 3.**
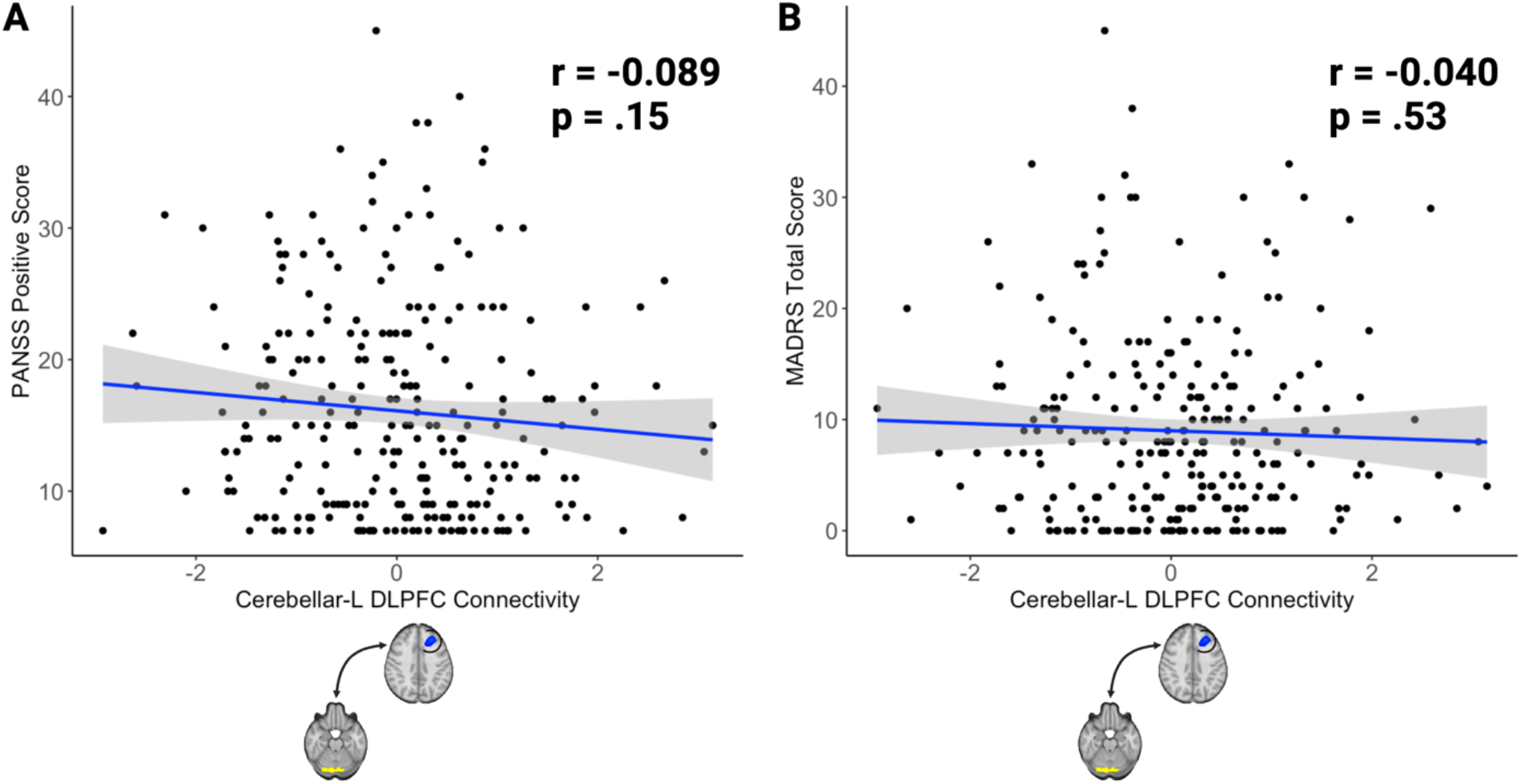
Cerebellar-Prefrontal Connectivity is Not Associated with Positive Symptom Severity or Depressive Symptoms. We observed that higher cerebellar-prefrontal connectivity was associated with lower negative symptom severity (Figure 2A & 2B, r =-0.17, p=.007). This connectivity relationship was specific to negative symptoms, as cerebellar-prefrontal connectivity was not associated with PANSS positive symptoms (r=-0.089, p=.15, Supplemental Figure 3A) or MADRS depressive symptoms (r=-0.040, p=.53, Supplemental Figure 3B).

